# Measuring protein conformational changes with genetically encoded bipyridylalanine and transition-metal ion FRET

**DOI:** 10.64898/2026.10.01.756074

**Authors:** Charlie DeFreest, Emerson Carmona, Sharona E. Gordon, William N. Zagotta

## Abstract

Transition-metal ion Förster resonance energy transfer (tmFRET) measures short-range distances and conformational changes in proteins, but its distance sensitivity depends on the donor– acceptor pair. Here, we establish the metal-chelating noncanonical amino acid bipyridylalanine (bpyA) as a flexible, genetically encoded acceptor platform whose Förster distance (R_0_) can be tuned without changing protein labeling. Cu²⁺, Cu²⁺ complexed with nitrilotriacetic acid (NTA), and Fe²⁺ form distinct acceptors at the same bpyA site, allowing distance sensitivity to be adjusted by changing the solution conditions. We incorporated bpyA at two sites in maltose binding protein (MBP) and labeled nearby engineered cysteines with the donor fluorophore BODIPY. The resulting [Cu(bpyA)]²⁺, [CuNTA(bpyA)]²⁺, and [Fe(bpyA)]²⁺ acceptors provided R_0_ values ranging from 13.9 to 21.2 Å. Analysis of time-resolved tmFRET data recovered donor–acceptor distance distributions and maltose-dependent changes consistent with structural predictions. At the 237/295 site pair, both copper and iron acceptors resolved the maltose-dependent decrease in distance. At the 309/322 site pair, [Cu(bpyA)]²⁺ did not appreciably resolve the distance increase, whereas adding NTA shifted R_0_ toward shorter distances and enabled detection of an increase from 12.1 to 14.4 Å. These results demonstrate that a single bpyA-labeled protein preparation supports multiple tmFRET acceptors, providing a cysteine-independent acceptor-labeling strategy with adjustable distance sensitivity.

## Introduction

Proteins are a group of dramatically diverse biological macromolecules which underlie virtually all biological processes in nature. Behaving as molecular machines, proteins perform a variety of functions, by undergoing dynamic changes in their three-dimensional structure (conformation) (3). Generally, these structural rearrangements occur in response to environmental stimuli, and, in turn, regulate the activity of the protein. Everything from metabolism, cell signaling, catalysis, transport and force generation are made possible by the dynamic nature of proteins.

Investigation of these protein structure-function relationships is hindered by our limited ability to probe the structural changes and energetics associated with conformational transitions (structural dynamics). X-ray crystallography and cryoEM provide high-resolution information on the static architecture of different conformations but little insight into the relative energies and transitions between conformational states. NMR and EPR spectroscopy can measure the dynamics of different regions of the protein, but provide less information about the precise structural changes. In addition, these approaches typically require purified protein, large sample quantities, and nonphysiological experimental conditions, and most NMR measurements are further restricted to relatively small proteins (<30 kDa).

Fluorescence Resonance Energy Transfer (FRET) provides an attractive approach to studying protein structural dynamics (4). FRET measurements can be conducted under physiological conditions, do not require purified protein, and can be performed at very low, even single molecule, concentrations. FRET involves non-radiative, distance-dependent energy transfer from an excited fluorescent donor to an absorbing acceptor (5). The efficiency of energy transfer (E) can be directly observed by the quenching of donor fluorescence and can be used to calculate the donor-acceptor distance. By site-specifically labeling a protein with donor and acceptor molecules, FRET can report distances between sites on the protein, as well as changes in distance that occur as the protein undergoes a conformational change.

Conventional FRET, however, presents important challenges for measuring intramolecular distances within proteins. FRET measured distances are most precisely determined near the Förster radius (R_0_), the donor-acceptor distance at which E = 0.5. Standard FRET experiments use a pair of fluorophores with overlapping emission and absorption spectra, which usually produces an R_0_ between 50 and 70 Å (Figure 1A, red) (5). This distance range is suitable for measuring distances between separate proteins or protein domains, but probing intradomain rearrangements requires access to shorter distances with higher spatial resolution (6,7). In addition, steady-state intensity-based FRET measurements can only measure a weighted-average distance (8). Since intramolecular distances in dynamic proteins are heterogeneous, arising from multiple conformations, these weighted average distances can be very different from the actual distances. Finally, the probes are generally introduced by cysteine labeling making it difficult to orthogonally label both the donor and acceptor on the same protein.

**Figure 1.**
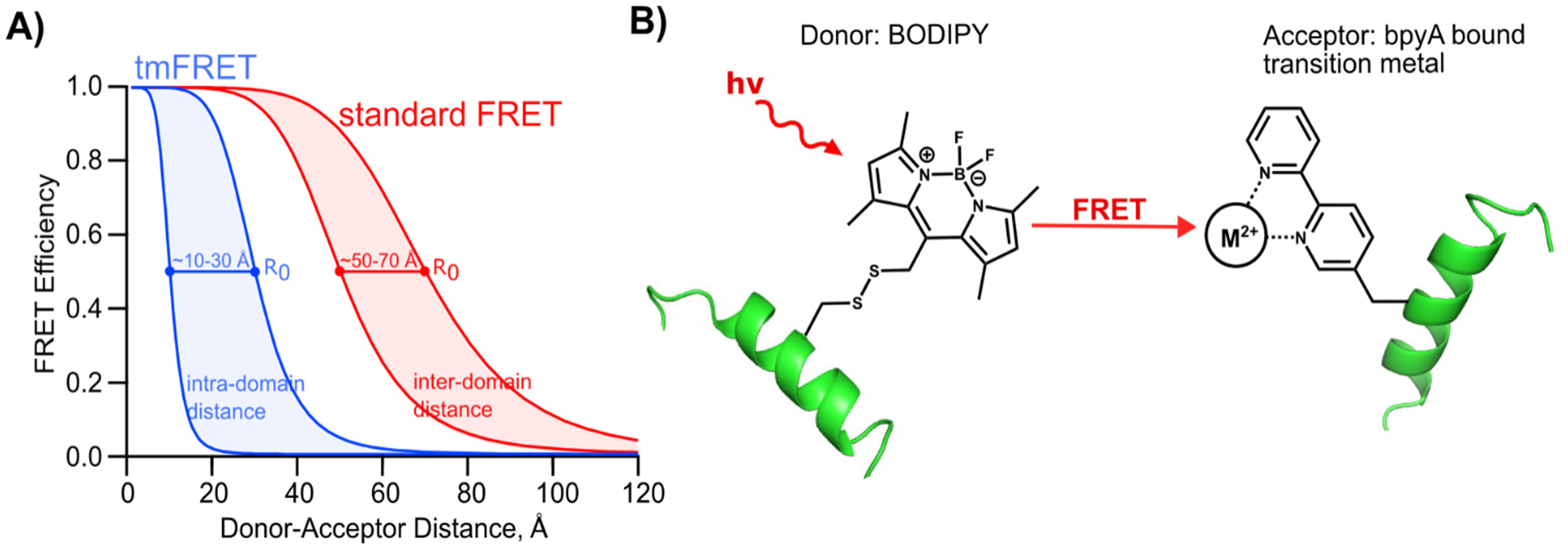
Distance dependence of transition-metal ion FRET (tmFRET). (A) FRET efficiency versus donor–acceptor distance (Förster curves) for tmFRET (blue) and conventional two-fluorophore FRET (red). The shorter Förster distances (R_0_) of tmFRET shift its distance sensitivity toward the range of intradomain distances. (B) Schematic of tmFRET between a BODIPY donor attached to a cysteine side chain and a transition-metal ion acceptor coordinated by bipyridylalanine (bpyA).

Transition-metal ion FRET (tmFRET) addresses these challenges of standard FRET. tmFRET is a variation of FRET in which the acceptor is a nonfluorescent transition metal ion such as Cu^2+^ or Fe^2+^ (Figure 1B) (6,7,9–14). Transition metals absorb light in the visible range but have much lower extinction coefficients than fluorophores. This results in shorter R_0_ values as well as steeper distance dependence (Figure 1A, blue). The metal ions can also be attached to the protein with shorter linkers than most fluorophores and therefore reduce the distance heterogeneity for tmFRET. And labeling with metal is orthogonal to cysteine labeling so the donor and acceptor can be site-specifically attached to the protein.

Previously, we have shown that a high-affinity metal binding site can be introduced into proteins by labeling a cysteine with a cysteine reactive metal chelator (e.g. TETAC, cyclenM, and phenM) (15–18). This strategy, however, is not possible if the protein of interest contains native cysteine residues required for function that cannot be mutated or selectively blocked. Additionally, this strategy prevents orthogonal labeling if both the donor and acceptor are introduced via cysteine labeling.

To expand the applicability of tmFRET to more proteins, we sought a method for labeling with cysteine-independent metal chelators. One such method is to use genetic code expansion (GCE; (19)) to introduce a noncanonical amino acid that can bind transition metal ions with high affinity. A chelator which meets these requirements is the non-canonical amino acid bipyridylalanine (bpyA) (Figure 1B). BpyA has high (nanomolar) affinity for binding transition metals (20) and has previously been introduced into proteins via GCE (21). Here, we used GCE to incorporate bpyA into a model protein, maltose binding protein (MBP). This system allowed us to orthogonally label the fluorescent donor and metal acceptor at different sites in the protein. Changing metal identity and coordination can tune the distance sensitivity of tmFRET (6,7,16). Here, we show that metal-bound bpyA can act as an acceptor for tmFRET and reveal distances and maltose-dependent changes in distance. We demonstrate the ability of bpyA to bind different metals and exhibit distinct coordination states, with different absorption spectra, thus providing a range of R_0_ values to tune tmFRET to the distances of interest. This approach expands the applicability of tmFRET to measure the structural dynamics of proteins that would be difficult to study with standard FRET methods.

## Results

### Incorporation of bpyA into MBP

To determine if bpyA is an effective metal-binding site for tmFRET, we used MBP as a model protein. MBP is a cysteineless clamshell-shaped protein with an equilibrium between its open and closed conformations regulated by the sugar maltose (Figure 2A). Binding maltose results in significant *closure* at the top of the clamshell, which is accompanied by a smaller *opening* at the bottom of the clamshell.

**Figure 2.**
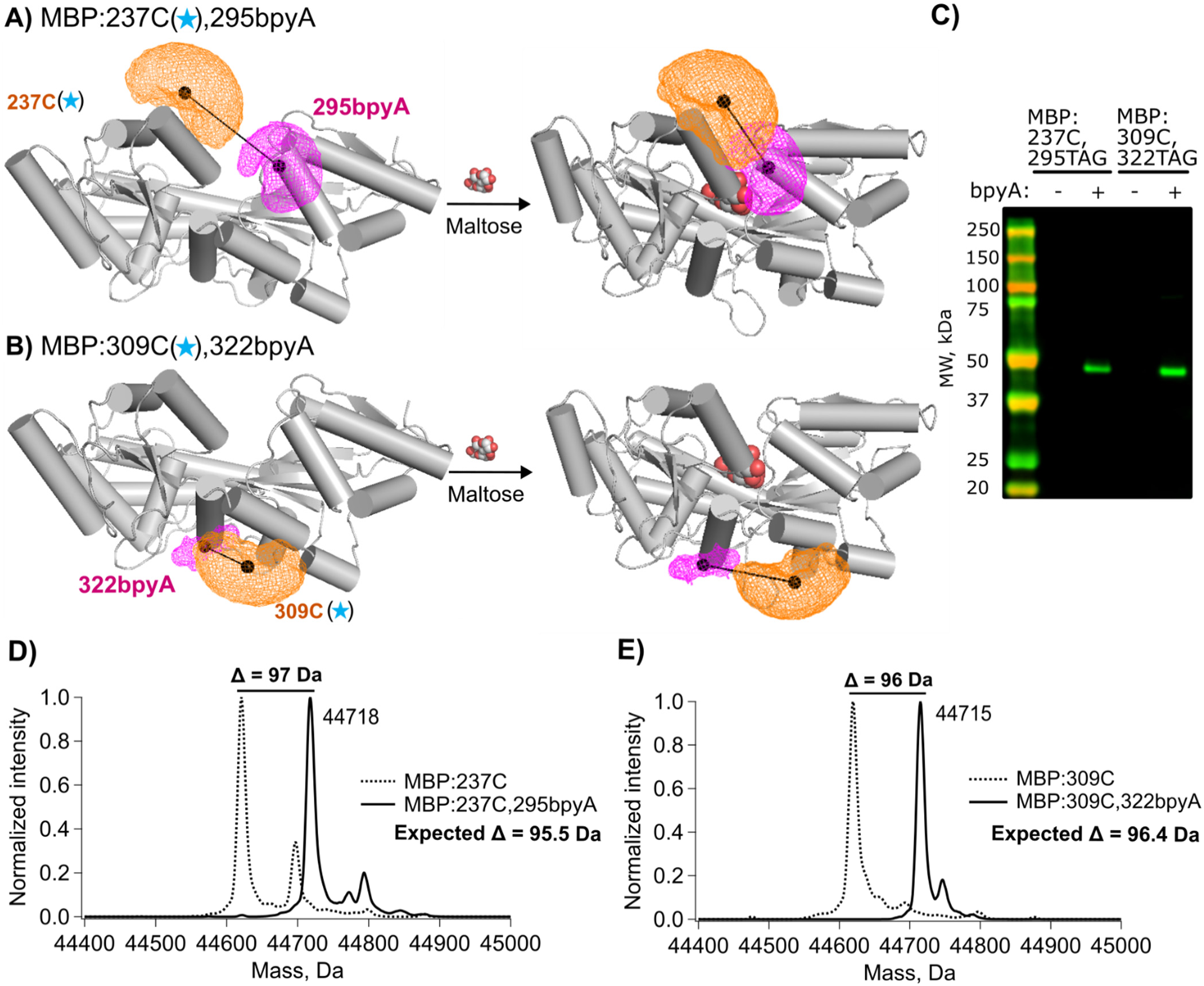
Site-specific incorporation of bpyA into maltose binding protein (MBP). (A, B) Rotamer distributions predicted by chiLife for BODIPY (orange) and Cu²⁺-bound bpyA (magenta) at sites 237 and 295 (A) or 309 and 322 (B). Distributions are shown on the crystal structures of apo MBP (left; PDB: 1OMP; (1)) and maltose-bound MBP (right; PDB: 1ANF; (2)). Black spheres mark the geometric centers of the distributions, and connecting lines indicate donor–acceptor distances. (C) Western blot of whole-cell lysates from E. coli expressing MBP with an amber codon at site 295 or 322 and the cognate aminoacyl-tRNA synthetase/tRNA pair, grown in the absence (−) or presence (+) of bpyA. Full-length MBP was detected using an antibody against its C-terminal twin-Strep tag. Molecular mass markers are shown at left. (D, E) Deconvoluted intact-protein mass spectra of MBP:237C,295bpyA (D) and MBP:309C,322bpyA (E) (solid curves), compared with the corresponding constructs lacking bpyA (dashed curves). Proteins were expressed in the presence of the genetic code expansion machinery and 500 µM bpyA. Observed mass shifts are indicated above the peaks; expected shifts are listed within each panel.

To assess the ability of tmFRET with metal-bound bpyA to report intramolecular distances and maltose-dependent increases *and* decreases in distance, we placed donor-acceptor site pairs across both the top and bottom of the clamshell. For the site-pair at the top of the clamshell, we genetically encoded bpyA at site 295 and introduced a single cysteine mutation at site 237 for modification with the donor fluorophore BODIPY-MTS (MBP:237C(★),295bpyA, Figure 2A). For the site-pair at the bottom of the clamshell, we encoded bpyA at site 322 and cysteine at site 309 (MBP:309C,322bpyA, Figure 2B). Control ‘donor-only’ constructs have the cysteine mutation at site 237 (MBP:237C) or 309 (MBP:309C) for labeling with the donor (★) but lack the bpyA metal-binding acceptor site. In the figure, the position of each label is depicted as a cloud representing the possible rotamers, which were modeled onto the X-ray crystal structures of MBP (grey) in the open (left, PDB: 1OMP; (1)) and closed (right, PDB:1ANF; (2)) conformations using the molecular modeling software package chiLife (22). At the top of the clamshell the donor-acceptor pair gets closer together with maltose (20.2 Å to 12.8 Å), and at the bottom of the clamshell, the donor-acceptor pair gets farther away with maltose (11.5 Å to 16.0 Å).

To site-specifically incorporate bpyA into sites 295 or 322 in MBP, we used genetic code expansion with amber-codon suppression. We co-expressed MBP constructs with the TAG stop codon at the desired site for bpyA and an aminoacyl tRNA synthetase and cognate tRNA previously developed for the incorporation of bpyA into proteins in *E. coli* (21). To validate the specific incorporation of bpyA at the engineered TAG sites, we performed Western blot analysis of crude cell lysates grown in the presence (+) or absence (−) of bpyA using an antibody against a C-terminal twin-Strep tag on MBP (Figure 2C). The presence of a band at the expected molecular weight for MBP-2x-Strep (∼45 kDa) only when the bpyA amino acid was supplied to the cells shows that only bpyA, and not cognate amino acids, was incorporated in the TAG sites.

To ensure that bpyA was added *specifically* at the TAG site, we used liquid chromatography-mass spectrometry (LC-MS). We performed LC-MS on the donor-only controls MBP:237C and MBP:309C, as well as the corresponding TAG site-containing constructs MBP:237C,295bpyA and MBP:309C,322bpyA. We found that the measured molecular weights of both donor-only controls matched the predicted values, whether or not we provided the cells with the GCE machinery and bpyA (Figure S1). In contrast, we observed shifts to higher molecular weights when a TAG site was present at sites 295 (Figure 2D, solid line) or 322 (Figure 2E, solid line), compared to the corresponding donor-only constructs (dashed lines). These observed molecular weight shifts were in agreement (within 2 Da) with the values expected for bpyA incorporation. Together, these results indicate that there was little to no off-target labeling occurring in the donor-only controls, and bpyA was site-specifically incorporated in MBP:237C(★),295bpyA and MBP:309C(★),322bpyA.

### 2,2-bipyridyl coordinates multiple types of transition metals or metal complexes yielding different R_0_s

The bpyA scaffold provides a flexible platform for tmFRET because it can coordinate multiple types of transition metals or metal complexes. This allows for distinct metal complexes with different absorption spectra and extinction coefficients, and consequently different R_0_ values, to be used in the same protein construct. To demonstrate the flexibility of the bpyA scaffold for tmFRET, we used three bpyA-acceptor species ([Cu(bpyA)]^2+^, [CuNTA(bpyA)]^2+^ and [Fe(bpyA)]^2+^) (Figure 3A). For each 1:1 [metal(bpyA)]^2+^ complex we measured the absorption spectrum in aqueous solution (Figure 3B). Using these spectra, we calculated the R_0_ value of each acceptor species with the donor used in this study (BODIPY) and plotted the FRET efficiency-distance relationship (Förster curves) for each pair in Figure 3C. The acceptor species [Cu(bpyA)]^2+^ has significant spectral overlap with the BODIPY emission spectrum, but with a low extinction coefficient (Figure 3B, cyan) resulting in an R_0_ value of 19.2 Å. The absorption spectrum of [CuNTA(bpyA)]^2+^ (Figure 3B, blue) is red-shifted relative to [Cu(bpyA)]^2+^, so there is less spectral overlap with BODIPY, resulting in a very short R_0_ value of 13.9 Å. The absorption spectrum of [Fe(bpyA)]^2+^ has a comparable range of overlap with BODIPY as [Cu(bpyA)]^2+^, but with a much higher extinction coefficient (Figure 3B, red), further increasing the R_0_ value to 21.2 Å.

**Figure 3.**
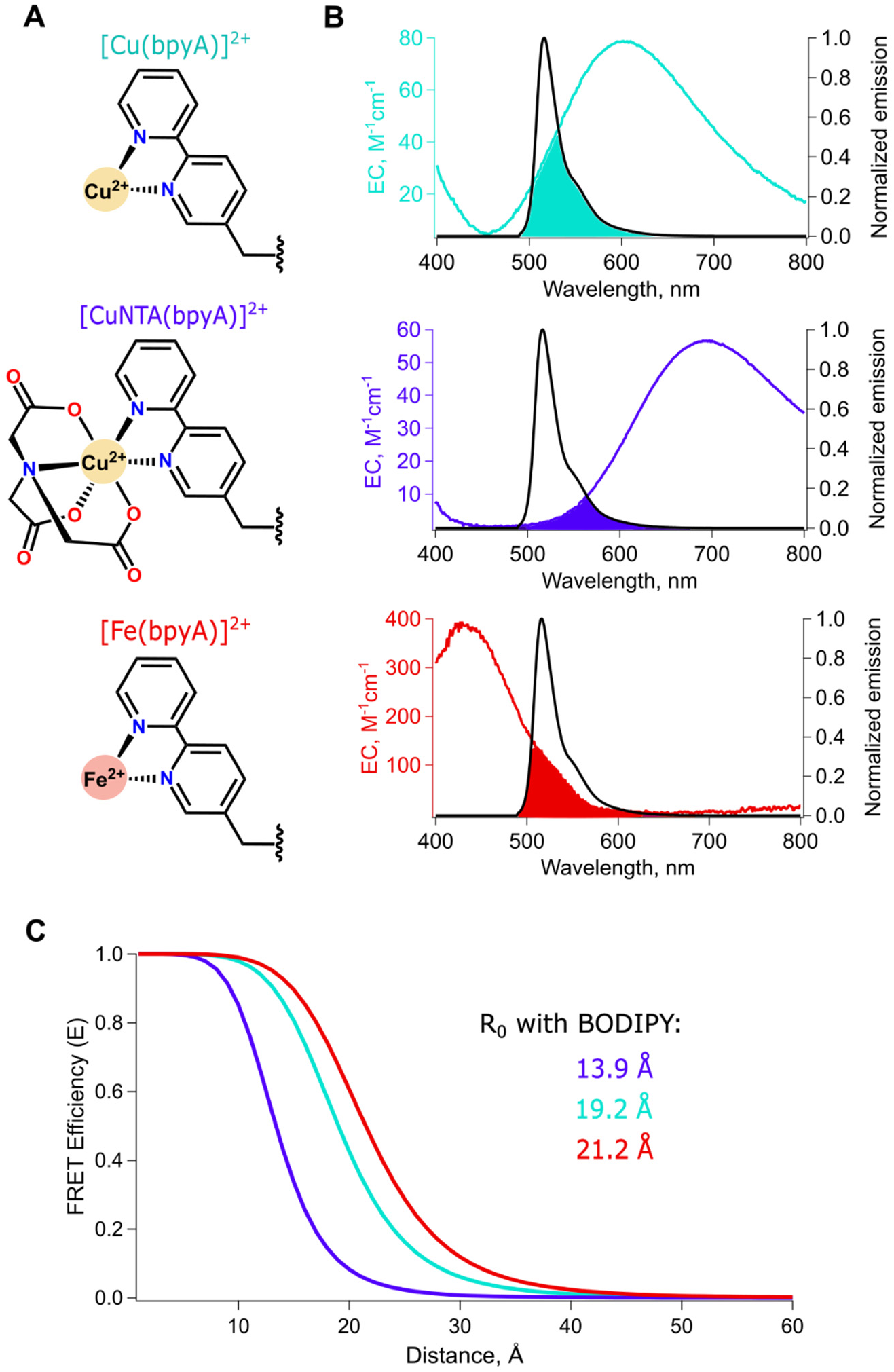
Spectral properties and distance dependence of bpyA-based tmFRET acceptors. (A) Structures of [Cu(bpyA)]²⁺ (cyan), [CuNTA(bpyA)]²⁺ (purple), and [Fe(bpyA)]²⁺ (red). Wavy bonds indicate connections to the protein backbone. (B) Absorption spectra of the three acceptors, expressed as molar extinction coefficients (colored curves; left axes), overlaid with the peak-normalized emission spectrum of BODIPY attached to MBP at site 237 (black curves; right axes). Shaded regions indicate spectral overlap. (C) FRET efficiency versus donor–acceptor distance for the three BODIPY–acceptor pairs. The corresponding R_0_ values are 19.2 Å for [Cu(bpyA)]²⁺, 13.9 Å for [CuNTA(bpyA)]²⁺, and 21.2 Å for [Fe(bpyA)]²⁺.

### Measuring tmFRET with fluorescence intensity quenching

Steady-state FRET can be measured as either acceptor sensitized emission or donor fluorescence quenching, both of which report the efficiency of energy transfer between donor and acceptor molecules. Because the transition-metal acceptors used in tmFRET are non-fluorescent, tmFRET is quantified directly from donor fluorescence quenching. Using the BODIPY-[Cu(bpyA)]^2+^ FRET pair, with its R_0_ of 19.2 Å, we measured tmFRET at the top of the MBP clamshell, where we expect a maltose-dependent decrease in donor-acceptor distance (see Figure 2A). In these experiments, we measured the steady-state fluorescence intensity of BODIPY initially and after the addition of a saturating concentration of metal (Figure 4). For MBP:237C(★),295bpyA, Cu^2+^ reversibly quenched the fluorescence intensity (Figure 4A,B). In contrast, for MBP:237C(★), without a bpyA, Cu^2+^ produced little to no quenching, indicating that quenching of MBP:237C(★),295bpyA resulted from specific [Cu(bpyA)]^2+^-mediated tmFRET. Importantly, tmFRET for MBP:237C(★),295[Cu(bpyA)]^2+^ was greater in the presence of maltose than in the absence of maltose. Together, these data demonstrate that bpyA functions as an effective genetically encodable tmFRET acceptor-binding site.

**Figure 4.**
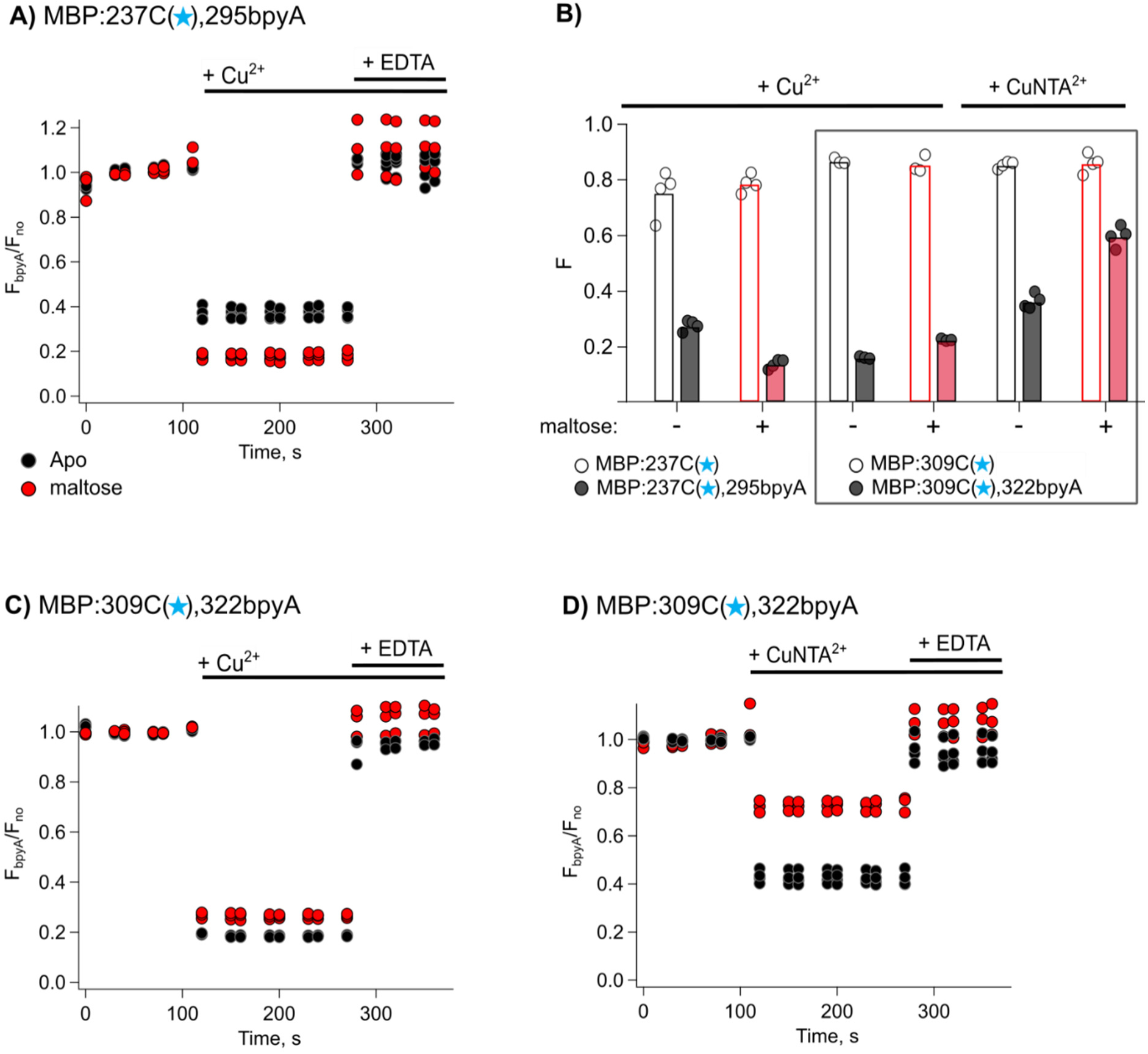
Steady-state tmFRET with bpyA-based copper acceptors in MBP. (A) Time courses of BODIPY fluorescence from MBP:237C(★),295bpyA following sequential addition of Cu²⁺ and EDTA, in the absence (black) and presence (red) of 2mM maltose. Fluorescence is normalized to its initial value and corrected by the fractional fluorescence of the corresponding donor-only construct. Bars indicate the periods following reagent addition. (B) Fractional fluorescence following addition of Cu²⁺ or CuNTA²⁺ for the indicated donor–acceptor constructs (filled symbols) and corresponding donor-only controls (open symbols), in the absence (black) and presence (red) of 2mM maltose. Individual measurements are shown overlaid on bars representing the average fractional fluorescence for each condition. (C, D) Time courses of normalized, donor-only-corrected fluorescence from MBP:309C(★),322bpyA following sequential addition of Cu²⁺ and EDTA (C) or CuNTA²⁺ and EDTA (D). Colors and reagent-addition bars are as in (A).

Unlike the site pair at the top of the clamshell, the site pair at the bottom of the clamshell (MBP:309C(★),322bpyA) undergoes a maltose-dependent *increase* in distance in the presence of maltose, and the distances are smaller (see Figure 2B). For MBP:309C(★),322bpyA, Cu^2+^ produced substantial fluorescence quenching, leaving little dynamic range to resolve the maltose-dependent change (Figure 4B, C). This result is consistent with the donor-acceptor distances in the absence and presence of maltose both being shorter than R_0_. A major advantage of tmFRET with bpyA is that we can tune the distance sensitivity simply by changing the metal complex. Shortening the R_0_ value using [CuNTA(bpyA)]²⁺ improved our ability to resolve the maltose-dependent distance change at the bottom of the clamshell (Figure 4B, D). These results demonstrate that modifying the coordinated metal complex extends the accessible distance range of tmFRET while using the same genetically encoded acceptor site.

### Measuring distance distributions in MBP:237C(★),265[Cu(bpyA)]^2+^ with time-resolved tmFRET

A more powerful way to quantify FRET is to measure fluorescence lifetimes—the latencies between photon absorption and emission (8,15,23). We refer to our application of this approach as time-resolved tmFRET. In the simplest cases, donor-only fluorescence lifetimes are described by a single exponential decay with a nanosecond time constant. The presence of a FRET acceptor accelerates the decay, reducing the lifetime, by introducing an additional distance-dependent nonradiative pathway. Heterogeneity in the donor-acceptor distances causes the donor lifetimes to become non-exponential, reflecting the weighted sum of the exponential lifetimes of many donor-acceptor distances (23). Fluorescence lifetime analysis can therefore be used to recover the underlying *distributions* of donor-acceptor distances, rather than a single weighted-average distance. Time-resolved tmFRET, therefore, captures nanosecond-long snapshots of the distance distributions, avoiding the averaging inherent in steady-state measurements. Additionally, lifetime measurements are largely independent of experimental variables such as fluorophore concentration, excitation intensity, and static quenching.

We performed time-resolved tmFRET to determine the distance distributions between 237C(★) and 295[Cu(bpyA)]^2+^ in the absence and presence of maltose. The fluorescence lifetimes of MBP:237C(★) (donor only) and MBP:237C(★),295bpyA (donor-acceptor) constructs in the presence of Cu^2+^ were measured with time-correlated single photon counting (TCSPC). As expected, the donor-only lifetimes were roughly single exponential (Figure 5A) and independent of Cu^2+^ (Figure S2). In contrast, in the presence of Cu^2+^, MBP:237C(★),295bpyA exhibited much shorter lifetimes consistent with Cu^2+^ binding to bpyA and acting as a FRET acceptor for 237C(★). Indeed, the lifetimes of MBP:237C(★),295[Cu(bpyA)]^2+^ were highly non-exponential, indicating a distribution of donor-acceptor distances contributing to the FRET (Figure 5A). We observed a further decrease in lifetimes upon the addition of a saturating concentration of maltose, reflecting the maltose-dependent closure of the clamshell that brings 237C(★) and 295[Cu(bpyA)]^2+^ closer together. Based on these results, we conclude that [Cu(bpyA)]^2+^ is a robust FRET acceptor for reporting donor-acceptor distance distributions and ligand-dependent changes in distance distributions.

**Figure 5.**
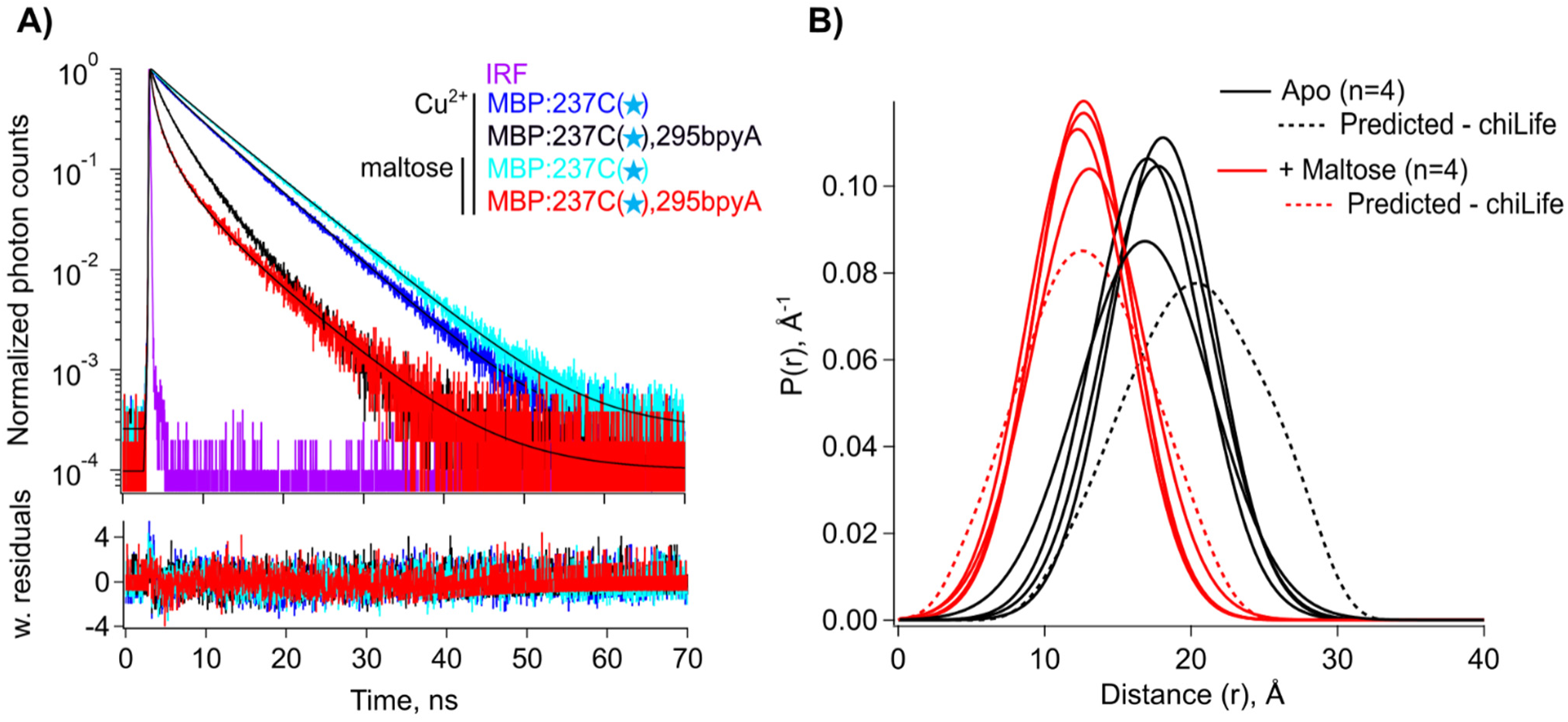
Time-resolved tmFRET of MBP:237C(★),2G5bpyA with [Cu(bpyA)]²⁺. (A) Representative fluorescence decays measured by time-correlated single-photon counting (TCSPC) in the presence of Cu²⁺, for MBP:237C(★),295bpyA without maltose (black) and with 2mM maltose (red), and the donor-only construct MBP:237C(★) without maltose (blue) and with 2mM maltose (cyan). Fits are overlaid on the decay traces, with weighted residuals shown below. The instrument response function (IRF) is shown in magenta. (B) Spaghetti plot of donor–acceptor distance distributions recovered from Gaussian-model fits to time-resolved tmFRET data without maltose (black solid curves; n = 4) and with maltose (red solid curves; n = 4), compared with distributions predicted by chiLife (dashed curves).

To recover donor-acceptor distance distributions from the TCSPC lifetimes, we used our previously developed model that assumes Gaussian distance distributions with the average distance, *r̅*, and standard deviation of the distribution (c) as free parameters (15). Fitting the data in Figure 5A with the Gaussian model revealed a maltose-dependent shift in *r̅* from 17.5 Å to 12.6 Å (solid lines, Figure 5B), consistent with chiLife predictions (dashed lines, Figure 5B). Thus, fits of time-resolved tmFRET data demonstrate that [Cu(bpyA)]^2+^ accurately reports the distance distributions associated with closure at the top of the MBP clamshell.

### Time-resolved tmFRET with MBP:306C(★),322bpyA

We next examined the site pair at the bottom of the clamshell, where maltose increases the donor-acceptor distance. We measured fluorescence lifetimes of MBP:309C(★) and MBP:309C(★),322bpyA, in the absence and presence of maltose (Figure 6A). In the presence of Cu^2+^, the lifetimes of MBP:309C(★),322bpyA were substantially shorter than the MBP:309C(★) lifetimes, confirming FRET occurred between the BODIPY donor and [Cu(bpyA)]^2+^ acceptor. However, the lifetimes were not appreciably longer in the presence of maltose, indicating that this FRET pair could not resolve the maltose-dependent change in distance at the bottom of the MBP clamshell (Figure 6B). This is not surprising because the donor-acceptor distances in the apo (11.5 Å) and maltose-bound (16.0 Å) states are both significantly shorter than the 19.2 Å R_0_ of BODIPY-[Cu(bpyA)]^2+^.

**Figure 6.**
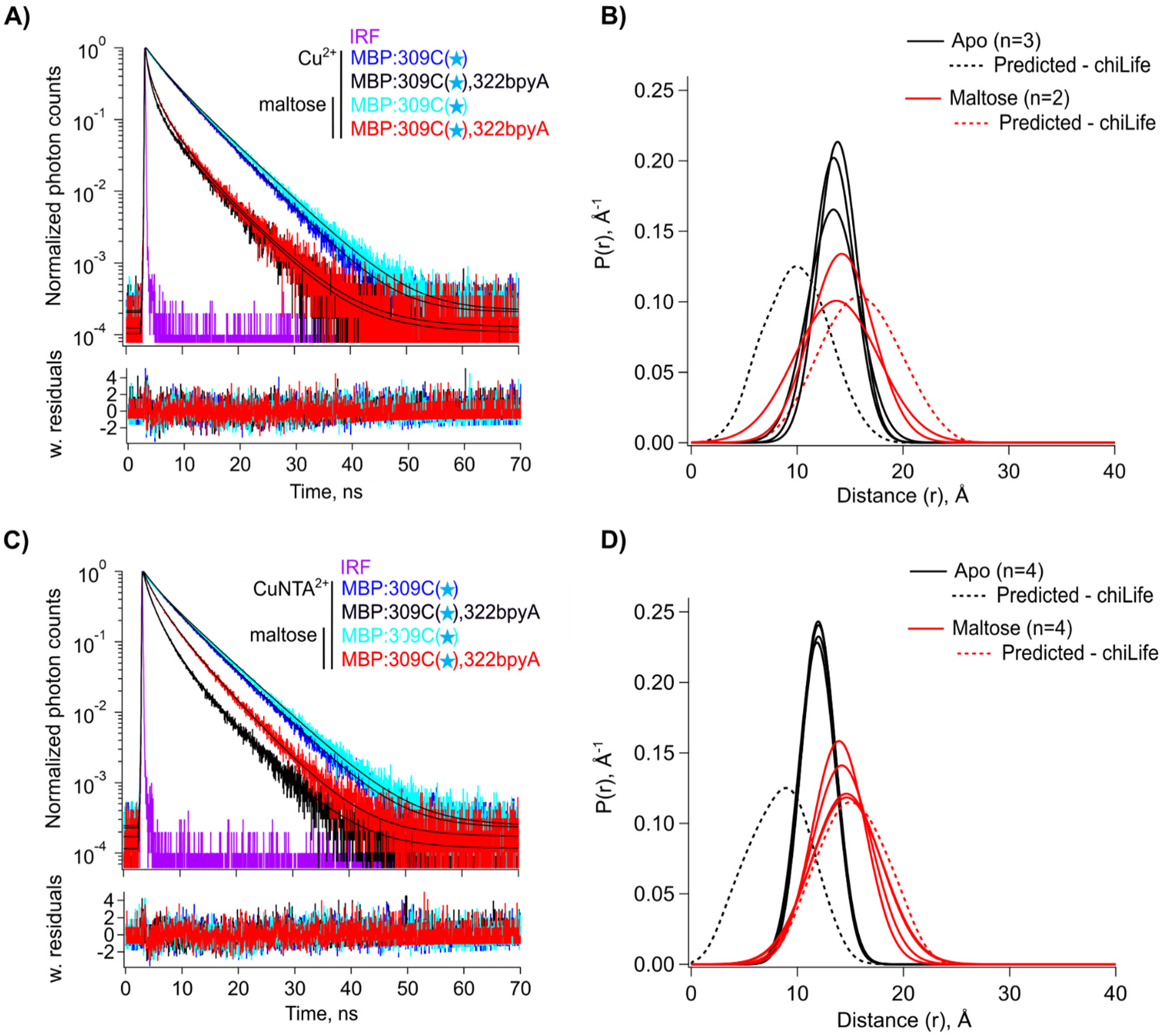
Time-resolved tmFRET of MBP:30GC(★),322bpyA with [Cu(bpyA)]²⁺ and [CuNTA(bpyA)]²⁺. (A, C) Representative TCSPC fluorescence decays in the presence of Cu²⁺ (A) or CuNTA²⁺ (C), for MBP:309C(★),322bpyA without maltose (black) and with 2mM maltose (red), and the donor-only construct MBP:309C(★) without maltose (blue) and with 2mM maltose (cyan). Fits are overlaid on the decay traces, with weighted residuals shown below. The IRF is shown in magenta. (B, D) Spaghetti plots of donor–acceptor distance distributions recovered from Gaussian-model fits using [Cu(bpyA)]²⁺ (B; n = 3 without maltose and n = 2 with maltose) or [CuNTA(bpyA)]²⁺ (D; n = 4 for each condition). Measured distributions without maltose (black solid curves) and with 2mM maltose (red solid curves) are compared with distributions predicted by chiLife (dashed curves).

To shift the distance sensitivity of tmFRET toward the shorter distances expected for MBP:309C(★),322bpyA, we used [CuNTA(bpyA)]^2+^ as an acceptor, which has a shorter R_0_ with BODIPY (13.9 Å). The donor-only lifetimes of MBP:309C(★) were independent of CuNTA^2+^ (Figure S3). In contrast to [Cu(bpyA)]^2+^, [CuNTA(bpyA)]^2+^ yielded a clear difference in lifetimes in the absence and presence of maltose (Figure 6C). Fits of the Gaussian model to the lifetimes recovered a maltose-dependent shift in *r̅* from 12.1 Å to 14.4 Å, consistent with the distances predicted by chiLife (Figure 6D). Thus, [CuNTA(bpyA)]^2+^ successfully resolved the maltose-dependent increase in distance in MBP:309C(★),322bpyA. Therefore, the ability to tune R_0_ by changing metal coordination allows the same genetically encoded bpyA site to be adapted to different distance regimes, extending the range of structural changes accessible to tmFRET without requiring different labeling sites or biochemical preparations.

### [Fe(bpyA)]^2+^ as a tmFRET acceptor to access longer distances

In addition to short distance measurements, the bpyA scaffold can be further utilized to probe longer distances. For example, Fe^2+^ bipyridyl complexes, which exhibit high extinction coefficients, can give much longer R_0_ values (16,24). We conducted time-resolved tmFRET experiments to evaluate [Fe(bpyA)]^2+^ as a tmFRET acceptor for resolving the maltose-dependent distance change in MBP:237C(★),295bpyA. Figure 7A shows the fluorescence lifetimes of MBP:237C(★) and MBP:237C(★),295bpyA with Fe^2+^ in the absence and presence of maltose. The fluorescence lifetime of MBP:237C(★) is unchanged by maltose (Figure 7A) or the addition of Fe^2+^ (Figure S4). However, there is a substantial maltose-dependent decrease in lifetimes in MBP:237C(★),295bpyA in the presence of Fe^2+^. Fits to the lifetimes recovered a maltose-dependent decrease in *r̅* from 17.7 Å to 13.2 Å, in agreement with chiLife predictions (Figure 7B). These results confirm that the BODIPY-[Fe(bpyA)]^2+^ donor-acceptor pair afforded accurate resolution of closure at the top of the clamshell, supporting bpyA as a scaffold for Fe^2+^-based acceptor design with extended R_0_ ranges.

**Figure 7.**
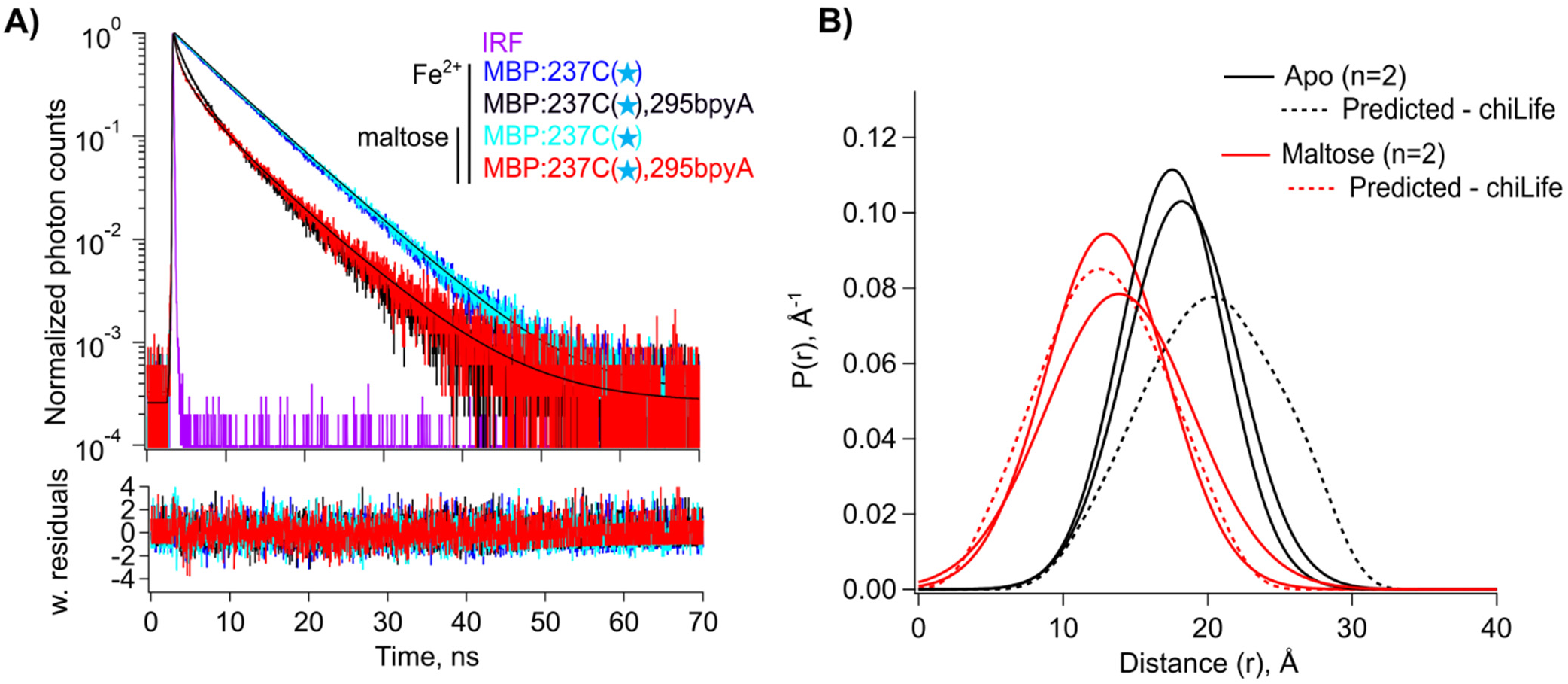
Time-resolved tmFRET of MBP:237C(★),2G5bpyA with [Fe(bpyA)]²⁺. (A) Representative TCSPC fluorescence decays in the presence of Fe²⁺, for MBP:237C(★),295bpyA without maltose (black) and in the presence of 2 mM maltose (red), and the donor-only construct MBP:237C(★) without maltose (blue) and with 2 mM maltose (cyan). Fits are overlaid on the decay traces, with weighted residuals shown below. The IRF is shown in magenta. (B) Spaghetti plot of donor– acceptor distance distributions recovered from Gaussian-model fits to time-resolved tmFRET data without maltose (black solid curves; n = 2) and with maltose (red solid curves; n = 2), compared with distributions predicted by chiLife (dashed curves).

## Discussion

In this work, we demonstrated that the noncanonical amino acid bpyA provides a versatile binding site for transition metal acceptors in tmFRET. BpyA was incorporated using GCE at two sites in maltose binding protein (MBP) and paired with nearby sites for incorporation of a donor fluorophore. In the presence of transition metal ions, tmFRET was observed both as quenching of donor fluorescence intensity and as a decrease in donor fluorescence lifetime. Analysis of time-resolved tmFRET revealed changes in the donor–acceptor distance distributions upon addition of maltose that were consistent with structural modeling. Moreover, different transition metal complexes produced different R_0_ values for the same protein construct, demonstrating that the distance range accessible by tmFRET can be tuned to the distances of interest. Together, these results establish the bpyA scaffold as a powerful platform for tmFRET measurements.

Previous tmFRET approaches have used engineered dihistidine sites (6,7,17,25–27) or cysteine-reactive metal chelators (15–17,24,28,29) to introduce transition-metal acceptors. BpyA provides several advantages over these approaches for measurements of protein structural dynamics. First, its compact metal-binding site provides a short linker for the acceptor, reducing distance heterogeneity arising from the rotameric distributions of the probes. Second, incorporation of bpyA by GCE allows the acceptor to be introduced in the presence of native cysteine residues and enables orthogonal labeling of the donor fluorophore, expanding the range of proteins accessible to tmFRET. Third, the high affinity of bipyridyl for transition metals (e.g., K_D_ = 8 nM for Cu^2+^; (20)) allows very low concentrations of metal to be used, reducing off-target metal binding and undesirable reactions of free metal ions, such as Cu^2+^-mediated oxidation of native cysteine residues. Finally, because bpyA can bind different transition metals and metal complexes, the distance range of tmFRET can be tuned without changing donor-acceptor sites or biochemical preparations.

The high metal affinity of bpyA also introduces several practical considerations. Because bpyA binds transition metals tightly, care must be taken to control which metal occupies the bpyA site, particularly in proteins containing native metal-binding sites where selective loading of different metals may be difficult. BpyA can also bind trace metals introduced during protein preparation or from solution. In our experiments, we therefore included EDTA in the measurement solution to chelate trace metals and added the transition metal of interest in excess of EDTA to ensure that the desired metal occupied the bpyA site. Metal-specific coordination chemistry must also be considered. For example, Fe^2+^ binds bipyridyl ligands cooperatively (20); care should be taken to prevent formation of [Fe(bpyA)_3_]^2+^ complexes during expression and purification.

Future work could extend bpyA-based tmFRET to fully orthogonal donor–acceptor labeling strategies that avoid cysteine chemistry altogether. For example, dual genetic code expansion could be used to introduce bpyA together with a second noncanonical amino acid carrying either a tetrazine or TCO group, enabling site-specific donor labeling through complementary tetrazine– TCO click chemistry (30,31). Such approaches could make tmFRET increasingly independent of native protein chemistry and broaden its application to more complex protein systems.

## Materials and Methods

### Reagents and Solutions

All chemicals were purchased from Millipore Sigma, unless otherwise specified. Potassium buffered Tris solution (KBT) was composed of 130 mM KCl (Sigma P3911-500G) and 30 mM Trizma base (Sigma, T6066; pH 7.5). <u>Auto-inducing medium (AIM) was composed of</u> 1.2 % w/v Tryptone, 2.4% w/v Yeast extract, 2 mM MgSO_4_, 0.5% w/v glycerol, 0.1% w/v glucose, 0.2% w/v a-Lactose, 25mM Na_2_HPO_4,_ 25mM KH_2_PO_4,_ 50mM NH_4_Cl 5mM Na_2_SO_4_, 50 μM FeCl_3_, 20 μM CaCl_2_, 10 μM MnCl_2_, 10 μM ZnSO_4_, 2 μM CoSO_4_, 2 μM CuCl_2_, 2 μM NiCl_2_, 2 μM Na_2_MoO_4_, 2 μM H_3_BO_3._

### Preparation of MBP-bpyA

Chemically competent BL-21 Gold (DE3) E.coli cells (Agilent, Cat#230132), were co-transformed with pETM11 containing MBP with C-terminal twin strep tag (16), and pEVOL containing the bpyA tRNA synthetase (derived from Mj Tyrosyl-RS, (MjTyrRS)) as well as the amber-suppressing tRNACUA (21)). Transformation was done via heat shock, exposing cells to 5 minutes on ice, followed by 30 seconds in a water bath at 42°C, followed by an additional 2 minutes on ice. Cells were then incubated with 1 mL SOC media at 37°C, 250rpm for 1 hr. Total volume of cells were then grown in 25 mL LB supplemented with 0.2% glucose, 50 ug/mL kanamycin, 34 ug/mL chloramphenicol at 37°C, 250 rpm O.N.

Pre-culture was diluted 1:100 in AIM supplemented with 100 ug/mL kanamycin, 34 ug/mL chloramphenicol, and grown at 37°C, 250 rpm until OD600 = 1.0. Once OD600 = 1.0, culture is supplemented with 0.1% L-arabinose to induce expression of tRNA synthetase and 500 µM bpyA. BpyA was dissolved in DMSO to a stock concentration of 100mM shortly before diluting in cell media. Cultures were then grown overnight at 20°C, 250 rpm.

Cells were harvested by centrifugation at 4,000 x g at 4°C for 15 minutes. Pellets were resuspended in KBT buffer supplemented with 1 mM EDTA, 1 mM PMSF, and 1mM benzamidine to a maximum concentration of 0.2 g pellet/mL KBT. Resuspended pellets are incubated with 2 mM beta mercaptoethanol and 0.5 mg/mL lysozyme for 30 minutes at 4°C with agitation, and then stored at −80°C until use.

Cell pellets were thawed in a water bath and supplemented with 1 mM PMSF, 1 mM benzamidine, 6 mM MgCl2, and 12.5 ug/mL DNase and incubated at 4°C with agitation for approximately 1 hour. 2 mM beta mercaptoethanol was added to the cell lysate, which was then sonicated 4 times for 1 min at 30% and 2.4 Output Power with a Branson Sonifier 450 in an ice bath with at least 1 min between cycles. Lysate was then spun in a JA-20 rotor at 40,000 x g for 30 minutes at 4C. The supernatant was harvested and passed over Streptactin XT gravity-flow resin (IBA Life Sciences) which had been equilibrated with KBT supplemented with 1 mM EDTA. Columns were then washed with at least 10 column volumes KBT, and protein was eluted using 50 mM biotin in KBT supplemented with 1 mM EDTA. 10% glycerol was added to the purified protein, which was aliquoted and flash frozen until use.

Purified protein was passed through a PD-10 column equilibrated with a low salt buffer (30 mM Tris, 10 mM KCl, pH = 7.5), injected to a HiTrap Q-HP anion exchange column with a 1 mL bed volume, and eluted with an increasing salt gradient (10 mM KCl to 1 M KCl) over 100 minutes. The fractions were then buffer exchanged using a PD-10 column equilibrated with KBT. In some cases, protein was not purified with anion exchange chromatography before use, with no difference in results.

### Labeling with BODIPY-MTS

Purified protein was reduced with 10 mM DTT, passed through a PD-10 column equilibrated with degassed KBT, and concentrated using a centrifugal concentrator. 1 mM EDTA was added to the sample followed by a 10x molar ratio of BODIPY-Methanethiosulfonate (BODIPY-MTS, Toronto Research Chemicals -B674875) dissolved in anhydrous DMSO. The reaction was incubated for 30 minutes on ice, an additional 10x molar ratio BODIPY-MTS was added, and the reaction was incubated for an additional 30 minutes. The sample was then passed through a PD-10 column before being injected to a Superdex 75 Increase 10/300 Size-Exclusion-Chromatography (SEC) column. Collected fractions were supplemented with 10% glycerol, and flash frozen in liquid nitrogen, and stored at −80°C.

### Mass-spectrometry

Determination of successful incorporation of bpyA was achieved using Liquid-Chromatography-Mass-Spectrometry (LC-MS) using a reversed phase HPLC column (Agilent BioHPLC PLRP-S Reversed-Phase column, 100Å, 2.1 x 50 mm, 3μm. Part#PL1912-1300) connected to an AB Sciex 5600 QTOF mass spectrometer. The mobile phase included 0.1% formic acid in MS-grade water or MS-grade acetonitrile. Data analysis was conducted using PeakView 2.2 software package.

### Transition-metal ions preparation

Cu^2+^ stock solution was prepared from chloride (CuCl_2_) salt at a concentration of 100 mM in water. CuNTA^2+^ stock was prepared outside the cuvette by mixing CuCl_2_ and NTA stocks in a 1:1.1 Cu^2+^:NTA molar ratio. Fe^2+^ stock solution was prepared from chloride (FeCl_2_) salt at a concentration of 200 mM in 1 M hydroxylamine (HA) solution in water (16). Cu^2+^ was added to cuvettes (for both fluorometry and TCSPC experiments) to a final concentration of 150 µM, CuNTA^2+^ was added at a final concentration of 1 mM, and Fe^2+^ was added at a final concentration of 350 µM. Given that 100 µM EDTA was present in the KBT sample buffer, there was approximately 50 µM free Cu^2+^, 900 µM free CuNTA^2+^, or 250 µM free Fe^2+^. For experiments involving Fe^2+^, 23 mM hydroxylamine hydrochloride was present in the KBT buffer solution which itself was prepared at pH =8.3.

### Fluorometry and spectrophotometry measurements

Fluorometry and spectrophotometry experiments were both performed with Starna (Atascadero, CA) sub-micro-fluorometer cells (100 μL). Absorption measurements were made using a Beckman Coulter DU 800 spectrophotometer (Brea, CA). Fluorometry experiments were performed using a Jobin Yvon Horiba FluoroMax-3 spectrofluorometer (Edison, NJ). We excited samples at 490 nm and recorded the emission at 516 nm at alternating 10 and 30 s intervals using the anti-photobleaching mode of the instrument (17). We used 5 nm slits for excitation and emission. Reagents (Cu^2+^, CuNTA^2+^, and EDTA) were added manually as 100-200 x stocks during the period between measurements by pipetting up and down in the cuvette without removing it from the instrument. For each experiment in the fluorometer, the background signal in the absence of protein was subtracted from the protein-containing signal. As described previously for time-course measurements (17), we normalized the fluorescence intensity values to the initial intensity signal.

### TCSPC lifetime measurements

Lifetime data were measured using a PicoQuant FluoTime 300 Fluorescence Lifetime Spectrometer (PicoQuant, Berlin, Germany) with 375 nm UV laser excitation and 516 nm emission, and single photon arrivals were recorded on a hybrid photomultiplier detector assembly (PMA-40). TCSPC data were acquired with the EasyTau2 measurement software (PicoQuant, Berlin, Germany). TCSPC data were analyzed with Igor Pro v9 (Wavemetrics) using custom scripts (https://github.com/zagotta/TDlifetime_program). The instrument response function was measured with Ludox.

### FRET model of time-domain fluorescence lifetime data to obtain Gaussian distance distributions

Fitting of the fluorescence lifetime data was as previously described (15). Briefly, the time-domain fluorescence lifetime data of the donor fluorophore in the presence of the acceptor were fit by model estimates for the time course of the decay, *Decay*(*t*). These fits were calculated from the convolution of the measured instrument response function, *IRF*(*t*) of the system with model estimates of the fluorescence lifetime of the donor in the presence of the acceptor, *I_DA_*(*t*) and buffer-only fluorescence *I_B_*(*t*):

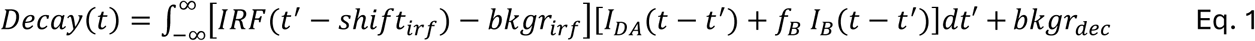

These variables are defined in Table 1.

**Table 1:**
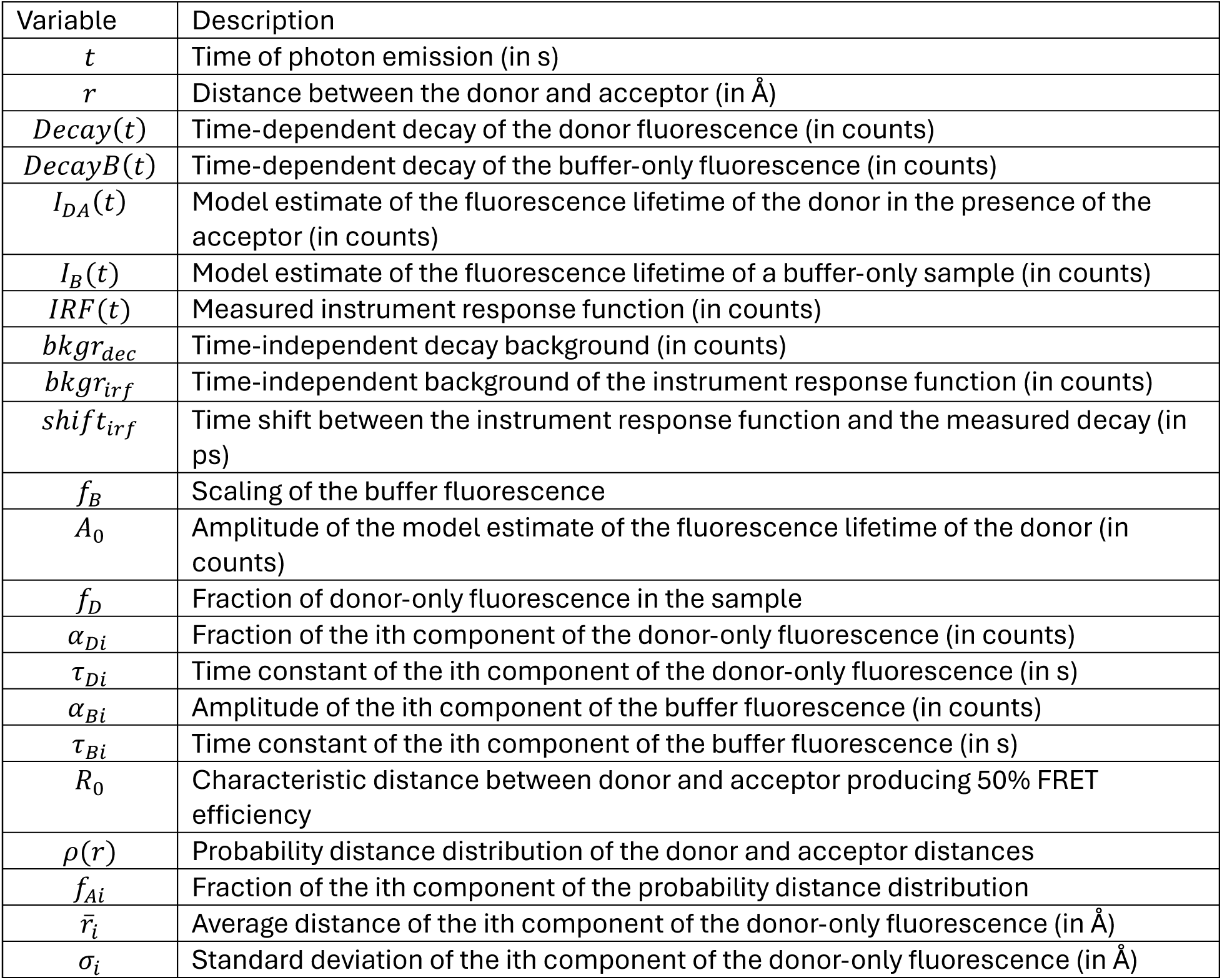

The model for fluorescence lifetime assumes a donor-only fluorescence lifetime with one or two exponential components, and FRET between a donor and acceptor separated by one or two Gaussian-distributed distances. This model predicts the following relationship for the fluorescence lifetime of the donor in the presence of acceptor, *I_DA_*(*t*):

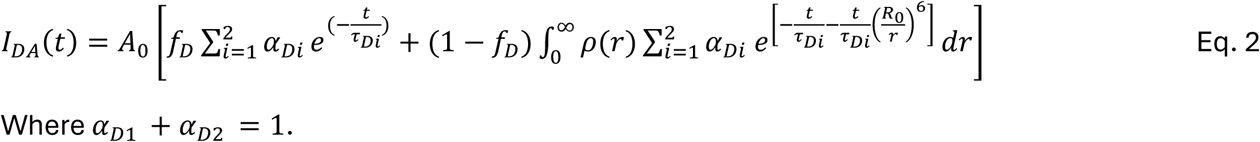

Where *α_D_*_1_ + *α_D_*_2_ = 1.

The density distribution of donor-acceptor distances, *ρ*(*r*), was assumed to be the sum of up to two Gaussians:

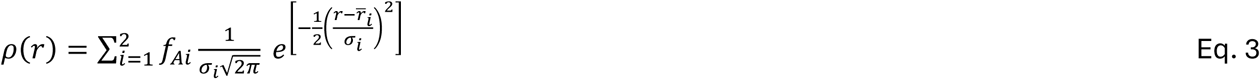

where *f_A_*_1_ + *f_A_*_2_ = 1.

The decay time course of the buffer-only sample, *DecayB*(*t*), was fit with a convolution of the measured instrument response function, *IRF*(*t*) with a multi-exponential model for the fluorescence lifetime, *I_B_*(*t*):

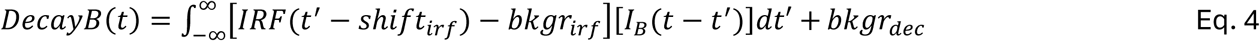

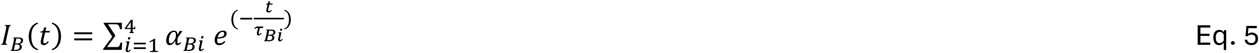

This model for the buffer-only fluorescence lifetime was then added to the model for the sample lifetime before convolving with the *IRF*(*t*) (Equation 1).

This FRET model for fluorescence lifetimes and FRET was implemented in Igor Pro v8 (Wavemetrics, Lake Oswego, OR) (code available at https://github.com/zagotta/TDlifetime_program). The FRET model was fit, with *χ*^2^ minimization, assuming a standard deviation equal to 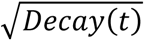, to individual decay time courses. There are 15 parameters in the parameter vector (*f_D_*, *τ_D_*_1_, *α_D_*_1_, *τ_D_*_2_, *R*_0_, *r̅*_1_, *σ*_1_, *f_A_*_2_, *r̅*_2_, *σ*_2_, *s*ℎ*ift_irf_*, *f_B_*, *A*_0_, *bkgr_dec_*, *bkgr_irf_*). *R*_0_ was fixed to a value previously determined from emission spectra of the donor and absorbance spectra of the acceptor using the Förster equation assuming *κ*^2^ = 2/3. *f_A_*_2_,was fixed to 0 or 1 for fits assuming a single Gaussian distance distribution. *f_B_* was fixed to the acquisition time for the experimental sample divided by the acquisition time for the buffer-only sample. And *bkgr_irf_*was always fixed to 0 as the average background photon count in the *IRF*(*t*) was <<1.

For the individual decay fits to the donor-only sample, *f_D_* was set to 1, and only *τ_D_*_1_, *α_D_*_1_, *τ_D_*_2_, *s*ℎ*ift_irf_*, *A*_0_, *bkgr_dec_* were allowed to vary. For the individual decay fits to the donor+acceptor samples, *τ_D_*_1_, *α_D_*_1_, *τ_D_*_2_were constrained to the previously determined values from the donor-only sample, and *f_D_*, *r̅*_1_, *σ*_1_, *f_A_*_2_, *r̅*_2_, *σ*_2_, *s*ℎ*ift_irf_*, *A*_0_, *bkgr_dec_* were varied.

## Acknowledgements

We thank the Oregon State University GCE4ALL (Center for Genetic Code Expansion for All) for their long-standing collaboration and providing bipyridylalanine. The bipyridylalanine aminoacyl tRNA synthetase plasmid was a kind gift of Peter G. Schultz (Scripps Research). We thank Drs. Max Tessmer and Stefan Stoll (Univesrsity of Washington) for technical assistance with chiLife. We also thank all members of the SEG and WNZ Laboratories for helpful conversations and support.

## References

1. Sharff, A. J., L. E. Rodseth, J. C. Spurlino, and F. A. Quiocho. 1992. Crystallographic evidence of a large ligand-induced hinge-twist motion between the two domains of the maltodextrin binding protein involved in active transport and chemotaxis. Biochemistry. 31(44):10657–10663, doi: 10.1021/bi00159a003, https://www.ncbi.nlm.nih.gov/pubmed/1420181.

2. Quiocho, F. A., J. C. Spurlino, and L. E. Rodseth. 1997. Extensive features of tight oligosaccharide binding revealed in high-resolution structures of the maltodextrin transport/chemosensory receptor. Structure. 5(8):997–1015, doi: 10.1016/s0969-2126(97)00253-0, https://www.ncbi.nlm.nih.gov/pubmed/9309217.

3. Henzler-Wildman, K., and D. Kern. 2007. Dynamic personalities of proteins. Nature. 450(7172):964–972, doi: 10.1038/nature06522, http://www.ncbi.nlm.nih.gov/pubmed/18075575.

4. Stryer, L., and R. P. Haugland. 1967. Energy transfer: a spectroscopic ruler. Proc Natl Acad Sci U S A. 58(2):719–726, http://www.ncbi.nlm.nih.gov/pubmed/5233469.

5. Lakowicz, J. R. 2006. Principles of fluorescence spectroscopy. Springer, New York.

6. Taraska, J. W., M. C. Puljung, and W. N. Zagotta. 2009. Short-distance probes for protein backbone structure based on energy transfer between bimane and transition metal ions. Proc Natl Acad Sci U S A. 106(38):16227–16232, doi: 10.1073/pnas.0905207106, http://www.ncbi.nlm.nih.gov/pubmed/19805285.

7. Taraska, J. W., M. C. Puljung, N. B. Olivier, G. E. Flynn, and W. N. Zagotta. 2009. Mapping the structure and conformational movements of proteins with transition metal ion FRET. Nat Methods. 6(7):532–537, doi: 10.1038/nmeth.1341, http://www.ncbi.nlm.nih.gov/pubmed/19525958.

8. Grinvald, A., E. Haas, and I. Z. Steinberg. 1972. Evaluation of the distribution of distances between energy donors and acceptors by fluorescence decay. Proc Natl Acad Sci U S A. 69(8):2273–2277, doi: 10.1073/pnas.69.8.2273, https://www.ncbi.nlm.nih.gov/pubmed/16592008.

9. Latt, S. A., D. S. Auld, and B. L. Vallee. 1972. Distance measurements at the active site of carboxypeptidase A during catalysis. Biochemistry. 11(16):3015–3022, http://www.ncbi.nlm.nih.gov/pubmed/5041907.

10. Latt, S. A., D. S. Auld, and B. L. Valee. 1970. Surveyor substrates: energy-transfer gauges of active center topography during catalysis. Proc Natl Acad Sci U S A. 67(3):1383–1389, http://www.ncbi.nlm.nih.gov/pubmed/5274465.

11. Horrocks, W. D., B. Holmquist, and B. L. Vallee. 1975. Energy transfer between terbium (III) and cobalt (II) in thermolysin: a new class of metal--metal distance probes. Proc Natl Acad Sci U S A. 72(12):4764–4768, http://www.ncbi.nlm.nih.gov/pubmed/1061067.

12. Richmond, T. A., T. T. Takahashi, R. Shimkhada, and J. Bernsdorf. 2000. Engineered metal binding sites on green fluorescence protein. Biochem Biophys Res Commun. 268(2):462–465, doi: 10.1006/bbrc.1999.1244, http://www.ncbi.nlm.nih.gov/pubmed/10679227.

13. Sandtner, W., F. Bezanilla, and A. M. Correa. 2007. In vivo measurement of intramolecular distances using genetically encoded reporters. Biophys J. 93(9):L45–47, doi: 10.1529/biophysj.107.119073, http://www.ncbi.nlm.nih.gov/pubmed/17766346.

14. Yu, X., X. Wu, G. A. Bermejo, B. R. Brooks, and J. W. Taraska. 2013. Accurate high-throughput structure mapping and prediction with transition metal ion FRET. Structure. 21(1):9–19, doi: 10.1016/j.str.2012.11.013, http://www.ncbi.nlm.nih.gov/pubmed/23273426.

15. Eggan, P., S. E. Gordon, and W. N. Zagotta. 2025. Domain coupling in allosteric regulation of SthK measured using time-resolved transition metal ion FRET. eLife. 14, doi: 10.7554/eLife.106892.

16. Gordon, S. E., E. G. B. Evans, S. C. Otto, M. H. Tessmer, K. D. Shaffer, M. T. Gordon, E. J. Petersson, S. Stoll, and W. N. Zagotta. 2024. Long-distance tmFRET using bipyridyl- and phenanthroline-based ligands. Biophys J. 123(14):2063–2075, doi: 10.1016/j.bpj.2024.01.034, https://www.ncbi.nlm.nih.gov/pubmed/38350449.

17. Zagotta, W. N., B. S. Sim, A. K. Nhim, M. M. Raza, E. G. Evans, Y. Venkatesh, C. M. Jones, R. A. Mehl, E. J. Petersson, and S. E. Gordon. 2021. An improved fluorescent noncanonical amino acid for measuring conformational distributions using time-resolved transition metal ion FRET. Elife. 10:e70236, doi: 10.7554/eLife.70236, https://www.ncbi.nlm.nih.gov/pubmed/34623258.

18. Gordon, S. E., M. Munari, and W. N. Zagotta. 2018. Visualizing conformational dynamics of proteins in solution and at the cell membrane. eLife. 7:e37248, doi: 10.7554/eLife.37248, https://www.ncbi.nlm.nih.gov/pubmed/29923827.

19. de la Torre, D., and J. W. Chin. 2021. Reprogramming the genetic code. Nat Rev Genet. 22(3):169–184, doi: 10.1038/s41576-020-00307-7, https://www.ncbi.nlm.nih.gov/pubmed/33318706.

20. Irving, H., and D. H. Mellor. 1962. The stability of metal complexes of 1,10-phenanthroline and its analogues. Part I. 1,10-Phenanthroline and 2,2’-bipyridyl. Journal of the Chemical Society (Resumed).(0):5222–5237, doi: 10.1039/JR9620005222, 10.1039/JR9620005222 (10.1039/JR9620005222).

21. Xie, J., W. Liu, and P. G. Schultz. 2007. A genetically encoded bidentate, metal-binding amino acid. Angew Chem Int Ed Engl. 46(48):9239–9242, doi: 10.1002/anie.200703397, https://www.ncbi.nlm.nih.gov/pubmed/17893898.

22. Tessmer, M. H., and S. Stoll. 2023. chiLife: An open-source Python package for in silico spin labeling and integrative protein modeling. PLoS Comput Biol. 19(3):e1010834, doi: 10.1371/journal.pcbi.1010834, https://www.ncbi.nlm.nih.gov/pubmed/37000838.

23. Haas, E., M. Wilchek, E. Katchalski-Katzir, and I. Z. Steinberg. 1975. Distribution of end-to-end distances of oligopeptides in solution as estimated by energy transfer. Proc Natl Acad Sci U S A. 72(5):1807–1811, doi: 10.1073/pnas.72.5.1807, https://www.ncbi.nlm.nih.gov/pubmed/1057171.

24. Zagotta, W. N., E. G. B. Evans, P. Eggan, M. H. Tessmer, K. D. Shaffer, E. J. Petersson, S. Stoll, and S. E. Gordon. 2024. Measuring conformational equilibria in allosteric proteins with time-resolved tmFRET. Biophys J. 123(14):2050–2062, doi: 10.1016/j.bpj.2024.01.033, https://www.ncbi.nlm.nih.gov/pubmed/38303511.

25. Dai, G., T. K. Aman, F. DiMaio, and W. N. Zagotta. 2019. The HCN channel voltage sensor undergoes a large downward motion during hyperpolarization. Nat Struct Mol Biol. 26(8):686–694, doi: 10.1038/s41594-019-0259-1, https://www.ncbi.nlm.nih.gov/pubmed/31285608.

26. Puljung, M. C., and W. N. Zagotta. 2013. A secondary structural transition in the C-helix promotes gating of cyclic nucleotide-regulated ion channels. J Biol Chem. 288(18):12944–12956, doi: 10.1074/jbc.M113.464123, http://www.ncbi.nlm.nih.gov/pubmed/23525108.

27. Dai, G., T. K. Aman, F. DiMaio, and W. N. Zagotta. 2021. Electromechanical coupling mechanism for activation and inactivation of an HCN channel. Nat Commun. 12(1):2802, doi: 10.1038/s41467-021-23062-7, https://www.ncbi.nlm.nih.gov/pubmed/33990563.

28. Hurwitz, S. M., W. N. Zagotta, S. E. Gordon, and S. Hoppins. 2026. Time-resolved tmFRET reveals GTP-coupled conformational changes in Mfn1. J Cell Biol. 225(7), doi: 10.1083/jcb.202511077, https://www.ncbi.nlm.nih.gov/pubmed/42080842.

29. Eggan, P., S. E. Gordon, and W. N. Zagotta. 2024. Ligand-coupled conformational changes in a cyclic nucleotide-gated ion channel revealed by time-resolved transition metal ion FRET. eLife. 13, doi: 10.7554/eLife.99854.

30. Koh, D. S., A. Stratiievska, S. Jana, S. C. Otto, T. M. Swanson, A. Nhim, S. Carlson, M. Raza, L. A. Naves, E. N. Senning, R. A. Mehl, and S. E. Gordon. 2024. Genetic code expansion, click chemistry, and light-activated PI3K reveal details of membrane protein trafficking downstream of receptor tyrosine kinases. eLife. 12, doi: 10.7554/eLife.91012, https://www.ncbi.nlm.nih.gov/pubmed/39162616.

31. Jana, S., E. G. B. Evans, H. S. Jang, S. Zhang, H. Zhang, A. Rajca, S. E. Gordon, W. N. Zagotta, S. Stoll, and R. A. Mehl. 2023. Ultrafast Bioorthogonal Spin-Labeling and Distance Measurements in Mammalian Cells Using Small, Genetically Encoded Tetrazine Amino Acids. J Am Chem Soc. 145:14608–14620, doi: 10.1021/jacs.3c00967, https://www.ncbi.nlm.nih.gov/pubmed/37364003.

