## Supplemental Figures for "Measuring protein conformational changes with genetically encoded bipyridylalanine and transition-metal ion FRET"

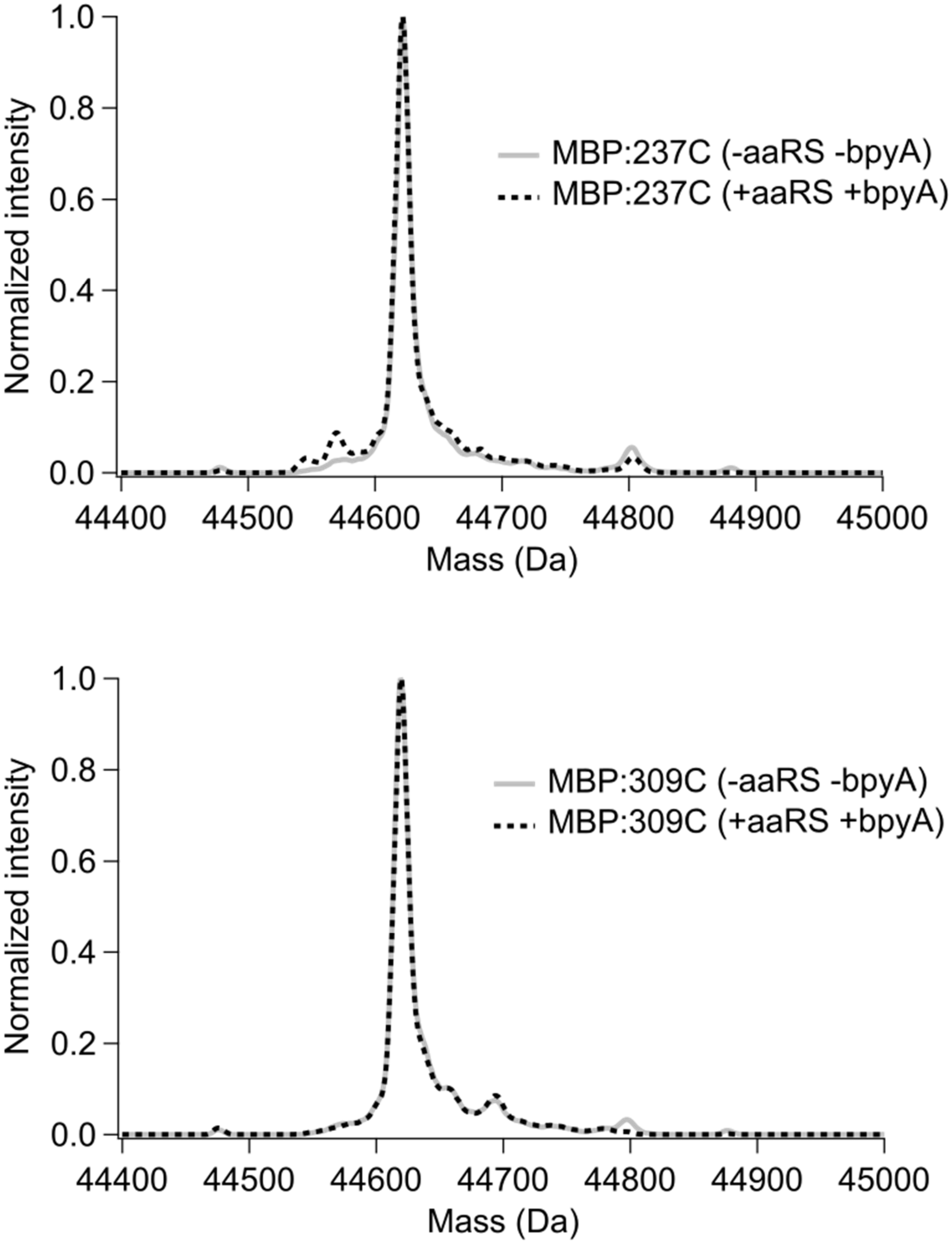


**Figure S1. Intact-protein mass spectra of donor-only MBP constructs expressed with and without the bpyA incorporation system.** Deconvoluted intact-protein LC-MS spectra of MBP:237C (top) and MBP:309C (bottom) expressed without the bpyA aminoacyl-tRNA synthetase and cognate suppressor tRNA and bpyA supplementation (gray solid curves) or with both (black dashed curves). Mass spectra were deconvoluted using PeakView software and normalized to their peak intensities.

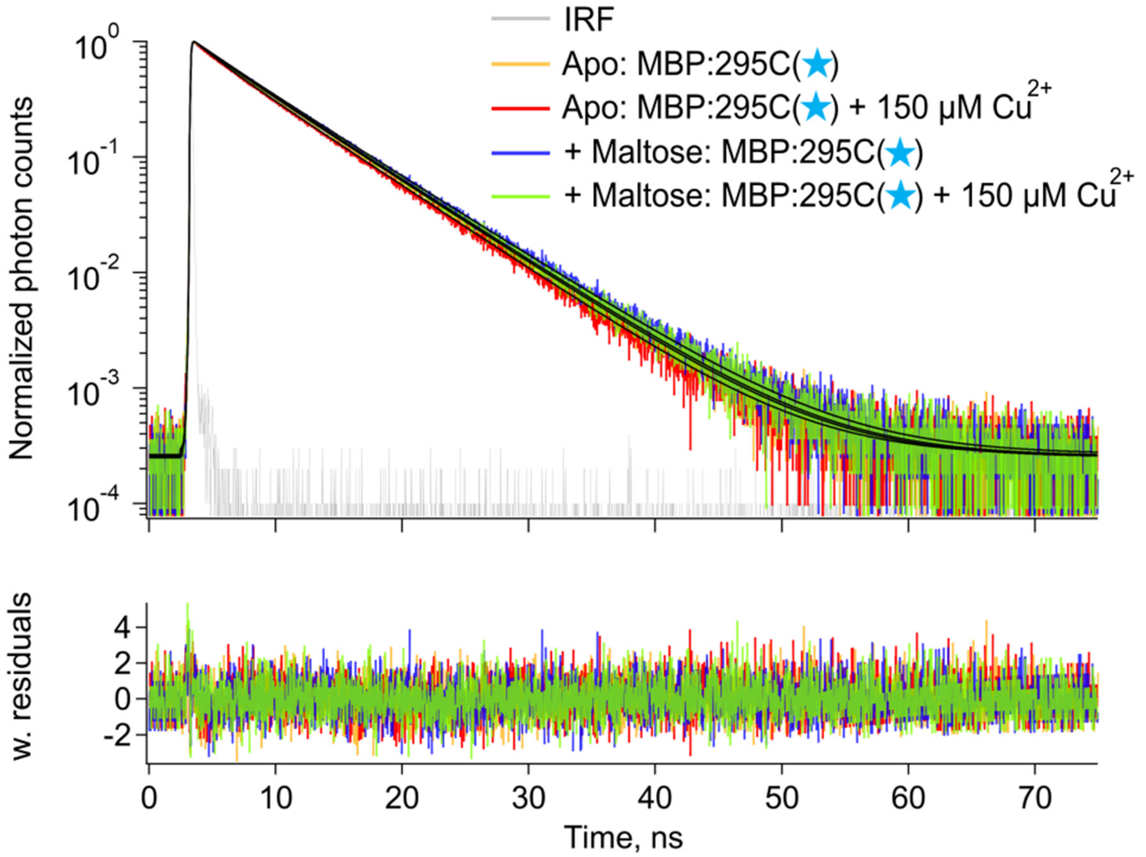


**Figure S2. Effect of $Cu^{2+}$ on donor-only MBP:237C(★) fluorescence decays.** Representative TCSPC fluorescence decays of BODIPY-labeled MBP:237C without maltose, before (orange) and after (red) addition of 150 µM $Cu^{2+}$, and with 2 mM maltose, before (blue) and after (green) addition of 150 µM $Cu^{2+}$. Fits are overlaid on the decay traces, with weighted residuals shown below. The instrument response function (IRF) is shown in gray.

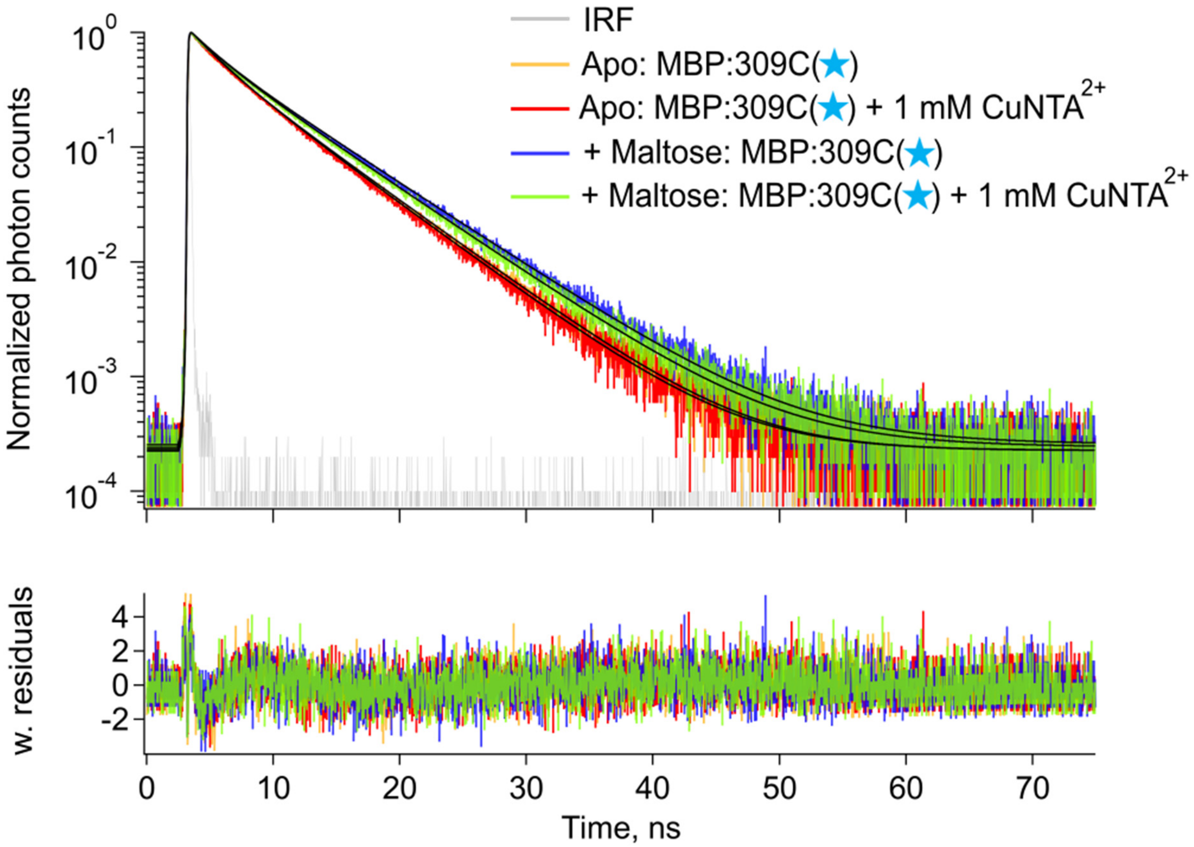


**Figure S3. Effect of $CuNTA^{2+}$ on donor-only MBP:309C(★) fluorescence decays.** Representative TCSPC fluorescence decays of BODIPY-labeled MBP:309C without maltose, before (orange) and after (red) addition of 1 mM $CuNTA^{2+}$, and with 2 mM maltose, before (blue) and after (green) addition of 1 mM $CuNTA^{2+}$. Fits are overlaid on the decay traces, with weighted residuals shown below. The IRF is shown in gray.

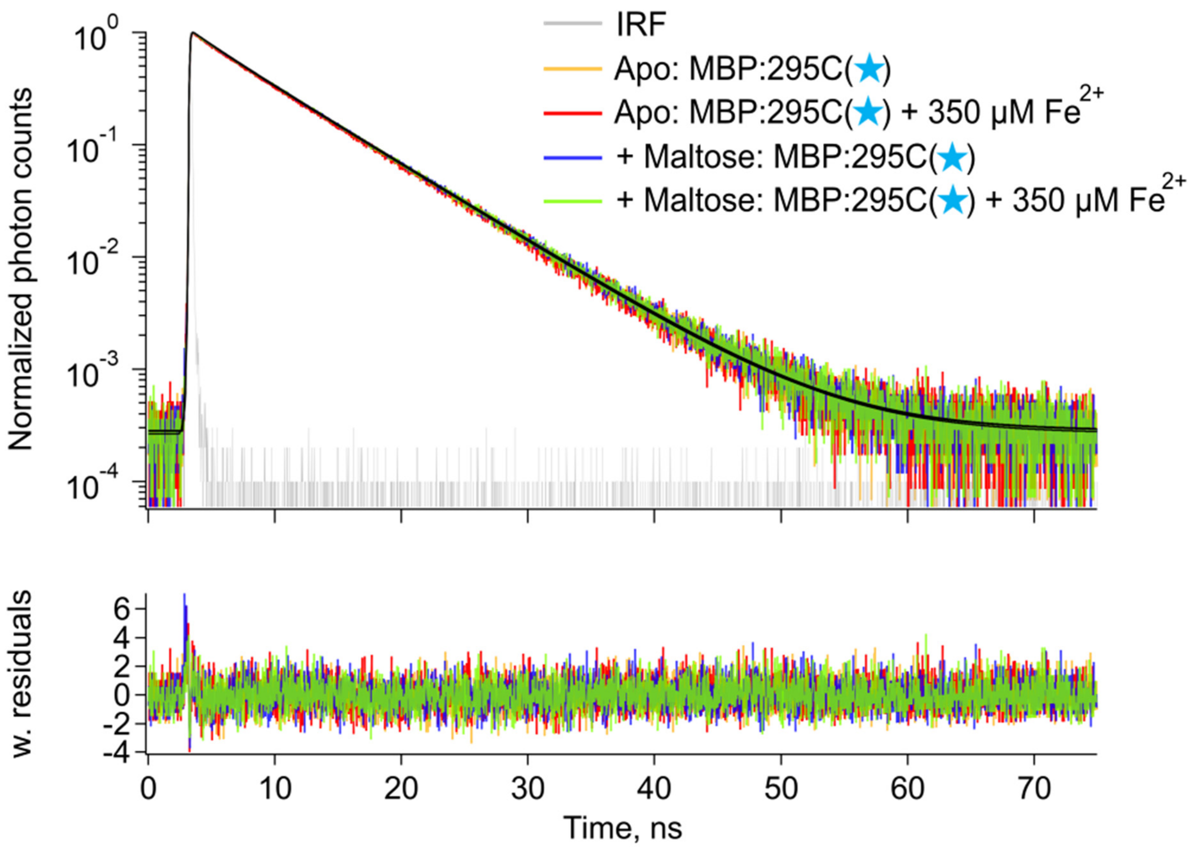


**Figure S4. Effect of $Fe^{2+}$ on donor-only MBP:237C(★) fluorescence decays.** Representative TCSPC fluorescence decays of BODIPY-labeled MBP:237C without maltose, before (orange) and after (red) addition of 350 µM $Fe^{2+}$, and with 2 mM maltose, before (blue) and after (green) addition of 350 µM $Fe^{2+}$. Fits are overlaid on the decay traces, with weighted residuals shown below. The IRF is shown in gray.
